# Overexpression of long noncoding RNA GAS5 suppresses tumorigenesis and development of gastric cancer by sponging miR-106a-5p through the Akt/mToR pathway

**DOI:** 10.1101/510511

**Authors:** Shuaijun Dong, Xiefu Zhang, Dechun Liu

## Abstract

Long non-coding RNAs (lncRNAs) have emerged as important regulators of human cancers. LncRNA GAS5 (GAS5) is identified tumor suppressor involved in several cancers. However, the roles of GAS5 and the mechanisms responsible for its functions in gastric cancer (GC) have not been well undocumented. Herein, the decreased GAS5 and increased miRNA-106a-5p levels were observed in GC and cell lines. GAS5 expression level was significantly inversely correlated with miRNA-106a-5p level in GC tissues. Moreover, luciferase reporter and qRT-PCR assays showed that GAS5 bound to miRNA-106a-5p and negatively regulated its expression in GC cells. Functional experiments showed that GAS5 overexpression suppressed GC cell proliferation, migration, and invasion capabilities and promoted apoptosis, while miRNA-106a-5p overexpression inversed the functional effects induced by GAS5 overexpression. *In vivo*, GAS5 overexpression inhibited tumor growth by negatively regulating miRNA-106a-5p expression. Mechanistic investigations revealed that GAS5 overexpression inactivating the Akt/mToR pathway by suppressing miRNA-106a-5p expression *in vitro* and *in vivo*. Taken together, our findings conclude the GAS5 overexpression suppresses tumorigenesis and development of gastric cancer by sponging miR-106a-5p through the Akt/mToR pathway.

## Introduction

Gastric cancer (GC), or stomach cancer, is a type of cancer that derives from the mucus-producing cells on the inside lining of the stomach and among that, the most common GC is adenocarcinoma. GC is the fifth most common cancer globally and the third most common cause of mortality associated with cancers (Pavlakis et al., 2016). Especially, Wang and his colleagues have reported that in China, it ranks second among cancer mortality, and its incidence still keeps rising (Wang et al., 2018). In spite of substantial improvements in surgical and chemotherapy treatments, the 5-year overall survival rate remains unsatisfactory by reason of diagnosis of the majority of patients with advanced GC accompanied by the relapse or metastasis (Luo et al., 2018). Furthermore, because of the intricate pathogenesis of GC, the molecular mechanisms of GC tumorigenesis and progression remain poorly understood. In recent years, research reported that GC develops through the accumulation of genetic and epigenetic alterations (Michigami et al., 2018). Therefore, it is important to disclose the underlying molecular mechanism of tumorigenesis to promote the development of new promising therapies for GC patients.

Recently, increasing studies have showed that noncoding RNAs (lncRNA) with limited or no protein-coding capacity are dysregulated in cancer progression (Deng et al., 2015, Wang et al., 2016). To date, many lncRNA expression alterations of multiple tumor-related genes have been reported to be associated with GC development. LncRNA maternally expressed gene 3 (MEG3) is was associated with metastatic GC, and its overexpression could constrain cell proliferation, migration, invasion and enhance cell apoptosis via sequestering oncogenic miR-181 in GC cells (S et al., 2015). Gastric adenocarcinoma predictive long intergenic noncoding RNA which is termed as GAPLINC were shown to contribute to the malignant phenotype, including the increased proliferation and invasiveness of GC cells, by activating CD44 expression, and inhibition of GAPLINC produced an significant reduction in tumor growth *in vivo* (S *et al*. 2015).

Growth arrest □ specific transcript 5 (GAS5) was firstly confirmed from mouse NIH 3T3 cells and identified as a possible tumor suppressor genes in many cancers (K et al., 2015). GAS5 has been reported to be aberrantly expressed in some cancers, including lung cancer (Wu et al., 2016), cervical cancer (W et al., 2014), breast cancer(GT 2016) and gastric cancer and function important roles in cell processes, such as proliferation, invasion, migration and apoptosis. Growing studies have disclosed the functional mechanism of lncRNAs in tumorigenesis by sponging specific miRNAs because of their direct interaction are attracting multi-field researchers’ attention (Ke et al., 2018, Toraih et al., 2018). However, the roles and potential mechnisms of GAS5 in GC are still not well clarified yet.

Here, bio-informatics analyses revealed a potential interaction between GAS5 and miRNA-106a-5p. Further we explored the expression profiles of GAS5 and miRNA-106a-5p in GC tissues and cell lines and analyzed their relationship. Moreover, functional and mechanism experiments were conducted *in vitro* and *in vivo*. Our findings provided a novel insight for the roles of GAS5 and miRNA-106a-5p in the development of GC and the regulatory mechanism involved.

## Results

### Decreased expression of GAS5 accompanied with increased miR-106a-5p in GC tissues and cells

Firstly, the expression levels of GAS5 and miRNA-106a-5p in patients with GC were determined by qRT-PCR. As shown in Fig 1A and B, GAS5 levels were significantly reduced in GC tissues, while the levels of miRNA-106a-5p were obviously higher than the adjacent normal tissues. And the analyses of GAS5 expression in different TNM stages showed that significant reduction of GAS5 levels in GC tissues at stage 3/4 was observed compared with stage 1/2 (Fig 1C). Then, the expression patterns of GAS5 and miRNA-106a-5p were compared by Pearson’s correlation coefficient and the presented results showed that the expression of GAS5 is negatively correlated with miRNA-106a-5p expression (Fig 1 D). In addition, the reduction of GAS5 levels (Fig 1 E) and the increase of miRNA-106a-5p levels (Fig 1 F) were also presented in GC cell lines (HCG-27 and SGC-7901) compared with human gastric epithelial cell line RGM-1. These results suggested that the aberrant expression of GAS5 and miRNA-106a-5p have a close relationship with GC development.

**Figure 1.**
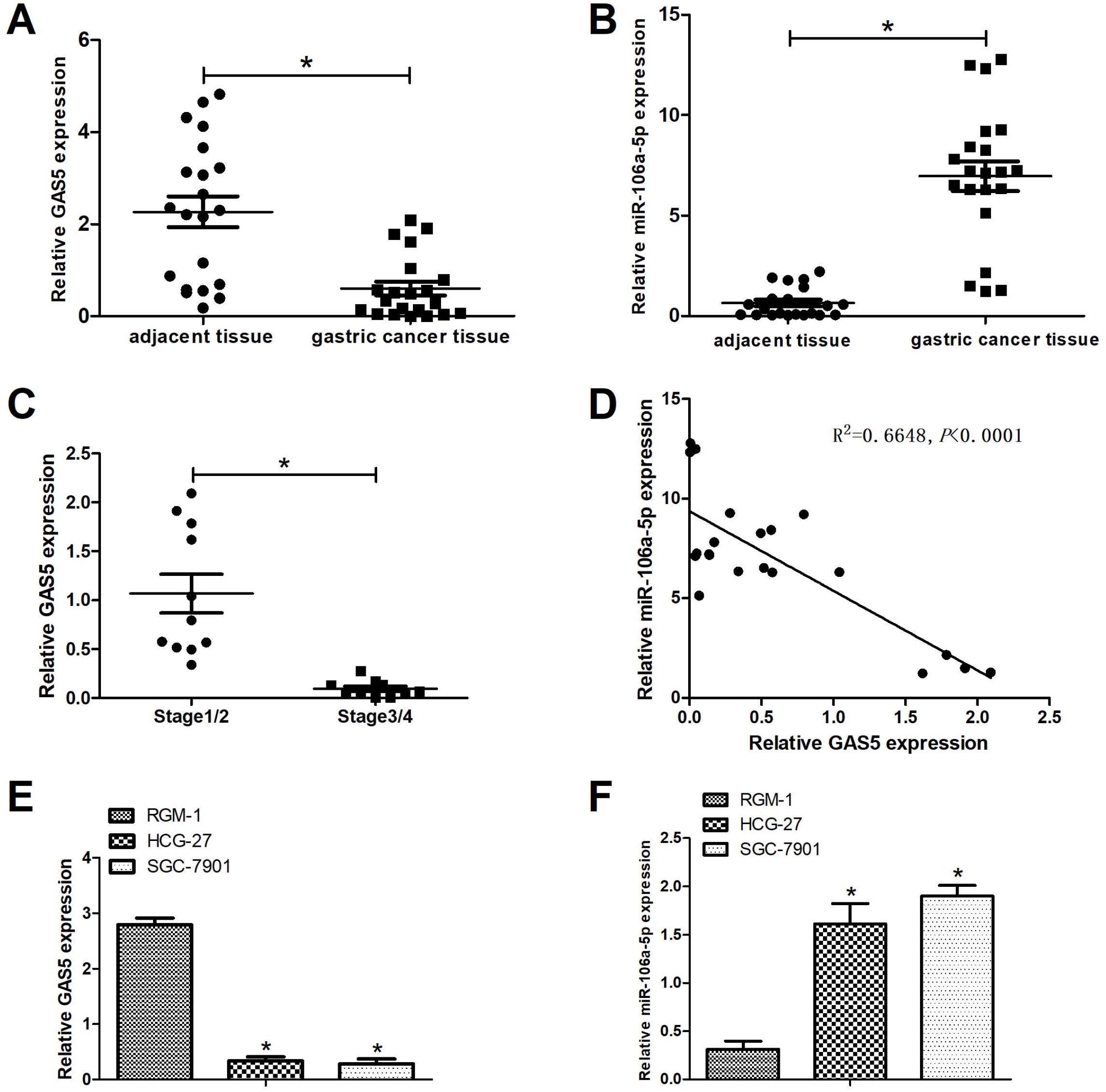
Decreased expression of GAS5 accompanied with increased miR-106a-5p in GC tissues and cells. Expression levels of GAS5 and miR-106a-5p were determined by qRT-PCR. The expression analyses of GAS5 (A) and miR-106a-5p (B) in GC and paired adjacent normal epithelial tissues from 21 GC patients. (C) The expression analyses of GAS5 in GC tissues at stages 1/2 and 3/4. (D) Pearson correlation between GAS5 and miR-106a-5p levels. The expression analyses of GAS5 (E) and miR-106a-5p (F) in Human gastric epithelial cell line RGM-1, GC cell lines (HCG-27 and SGC-7901). \**P* < 0.05.

### GAS5 interacted with miR-106a-5p and negatively regulated its expression in GC cells

Given the above results, we speculated the GAS5 could sponge to miRNA-106a-5p. Therefore, bioinformatics analyses were performed by software Megalign. As presented in Figure 2A, miRNA-106a-5p may be a candidate binding miRNA of GAS5. Further, the results from luciferase reporter assay showed that the luciferase activity from GAS5-WT was significantly decreased in miRNA-106a-5p transfected group. However, miRNA-106a-5p transfection has no significant effects on the luciferase activities of GAS5-MUT in 293T cells (Figure. 2C). Moreover, qRT-PCR analyses showed that GAS5 overexpression significantly in repressed the expression of miRNA-106a-5p in HCG-27 (Figure 2C) and SGC-7901 (Figure 2D) cells. Taken together, these data suggested that GAS5 directly targeted miR-106a-5p and negatively regulated its expression in GC cells.

**Figure 2.**
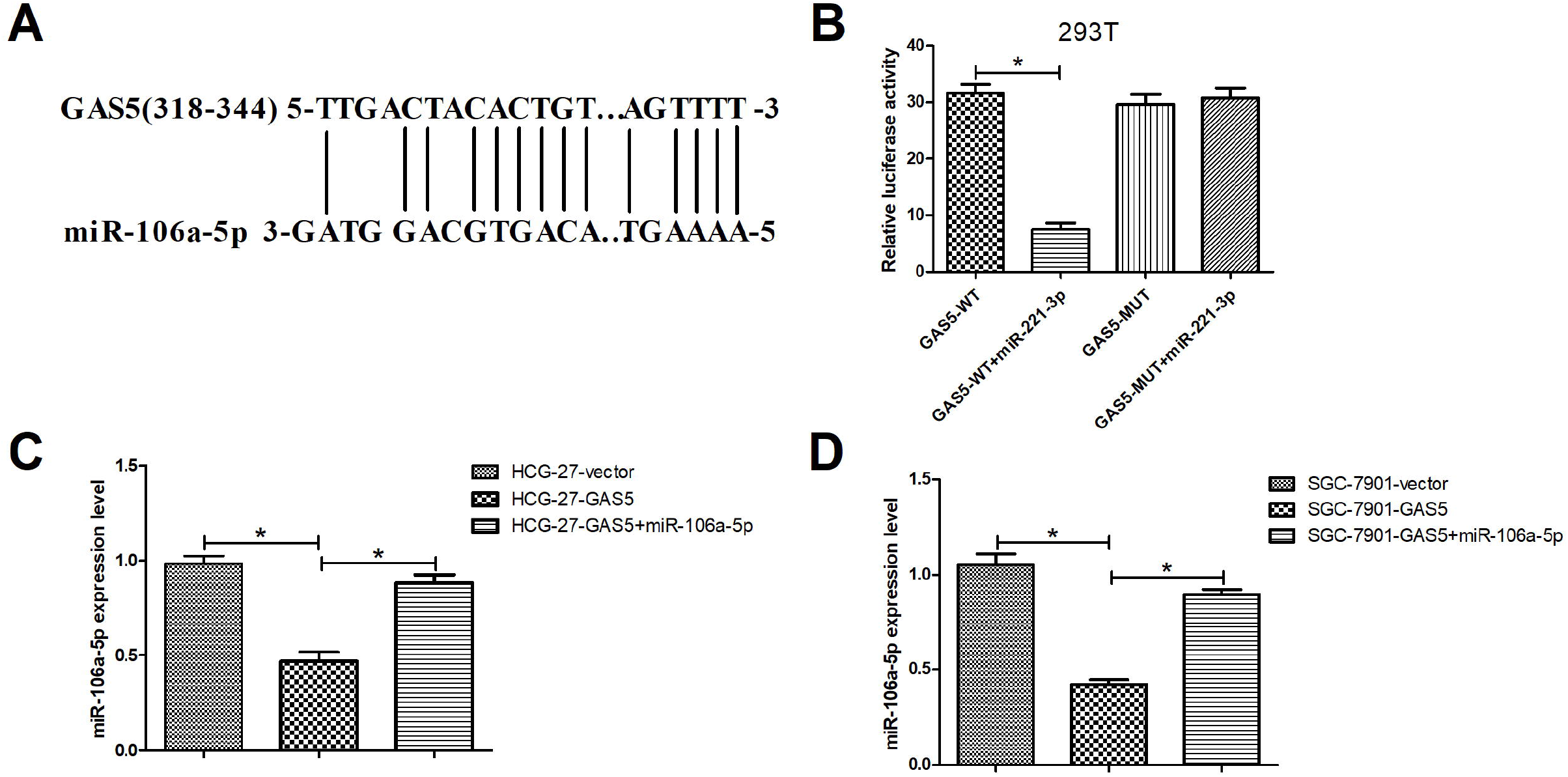
GAS5 interacted with miR-106a-5p and negatively regulated its expression in GC cells. (A) Bioinformatics analyses were conducted to predict the possible binding site of miR-106a-5p in GAS5. (B) Dual-luciferase reporter assay was performed to determine the interaction of GAS5 and miR-106a-5p in 293T cells at 48 h. qRT-PCR analyses of miR-106a-5p in the HCG-27 (C) and SGC-7901 (D) cells transfected with GAS5, GAS5 + miR-106a-5p, or vector. \**P* < 0.05.

### GAS5 overexpression inhibited proliferation and induced apoptosis via regulating miR-106a-5p in GC cells

To further explore the biological roles of GAS5 and miRNA-106a-5p, a series of functional experiments were carried out. As exhibited in Figure 3A and B, overexpression of GAS5 significantly constrained the proliferation of HCG-27 and SGC-7901 cells. In addition, GAS5 upregulation obviously promoted apoptosis in HCG-27 (Figure 3C) and SGC-7901 (Figure 3D) cells, whereas miRNA-106a-5p overexpression evidently inversed the effect of GAS5 upregulation. These data implied that GAS5 overexpression supressed proliferation and induced apoptosis via regulating miR-106a-5p in GC cells.

**Figure 3.**
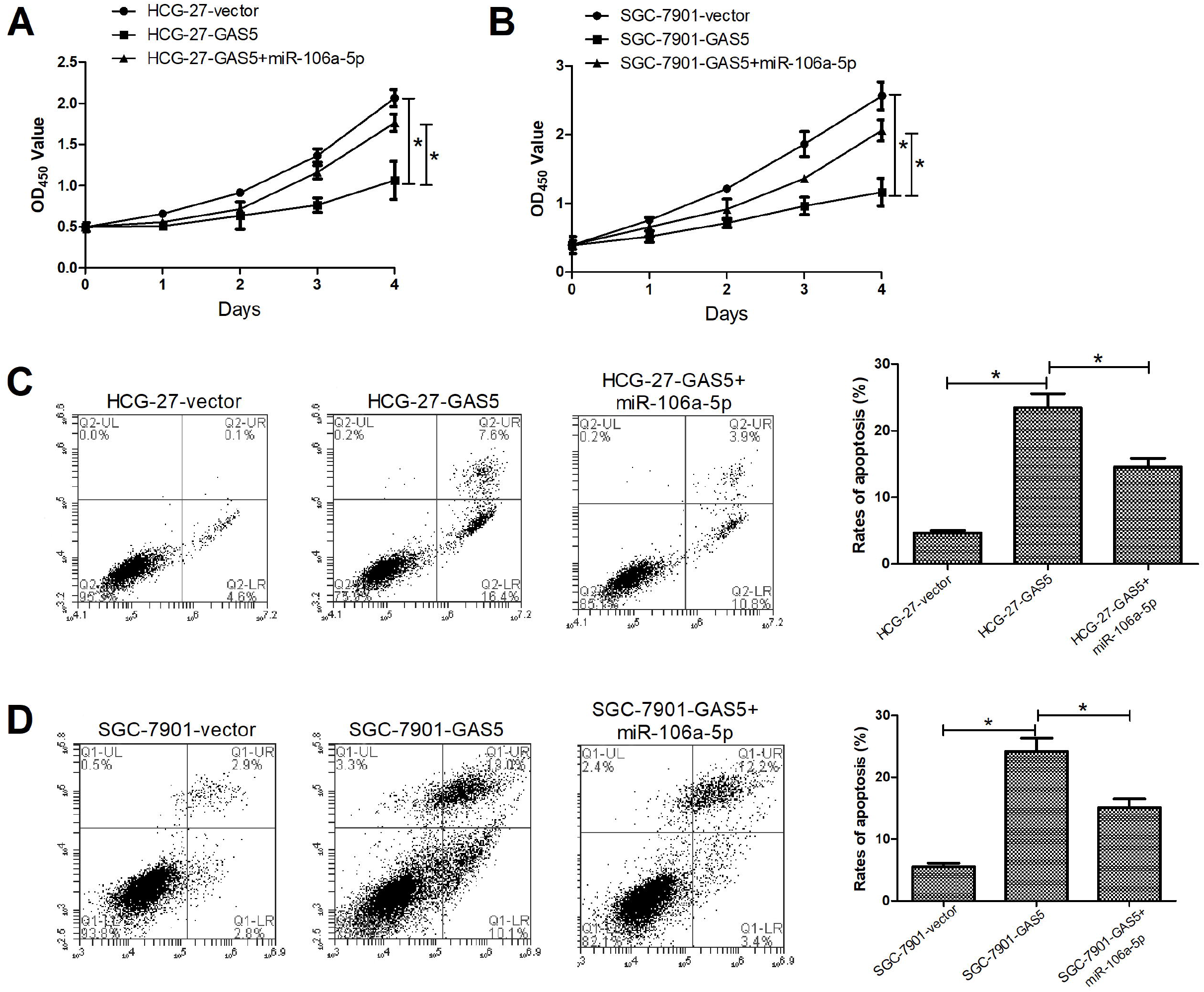
GAS5 overexpression inhibited proliferation and induced apoptosis via regulating miR-106a-5p in GC cells. CCK-8 assay were used to proliferation of HCG-27 (A) and SGC-7901 (B) cells transfected with GAS5, GAS5 + miR-106a-5p, or vector at indicated times (0, 1, 2, 3, and 4 days). Flow cytometry analyses of apoptosis rates of the transfected HCG-27 (C) and SGC-7901 (D) cells at 48 h. \**P* < 0.05.

### GAS5 overexpression repressed invasion and migration via regulating miR-106a-5p in GC cells

As shown in Figure A and B, transwell assays showed that upregulation of GAS5 significantly reduced the capability of GC cell invasion compared with control group, while the introduction of miRNA-106a-5p obviously weakened the repressive effects. Moreover, wound healing assay revealed that overexpression of GAS5 markedly constrained the migration of HCG-27 (Fig 4C) and SGC-7901 (Fig 4D) cells, and miRNA-106a-5p significantly inversed the suppressive effects of overexpression of GAS5. These results suggested that GAS5 overexpression repressed invasion and migration via regulating miR-106a-5p in GC cells.

**Figure 4.**
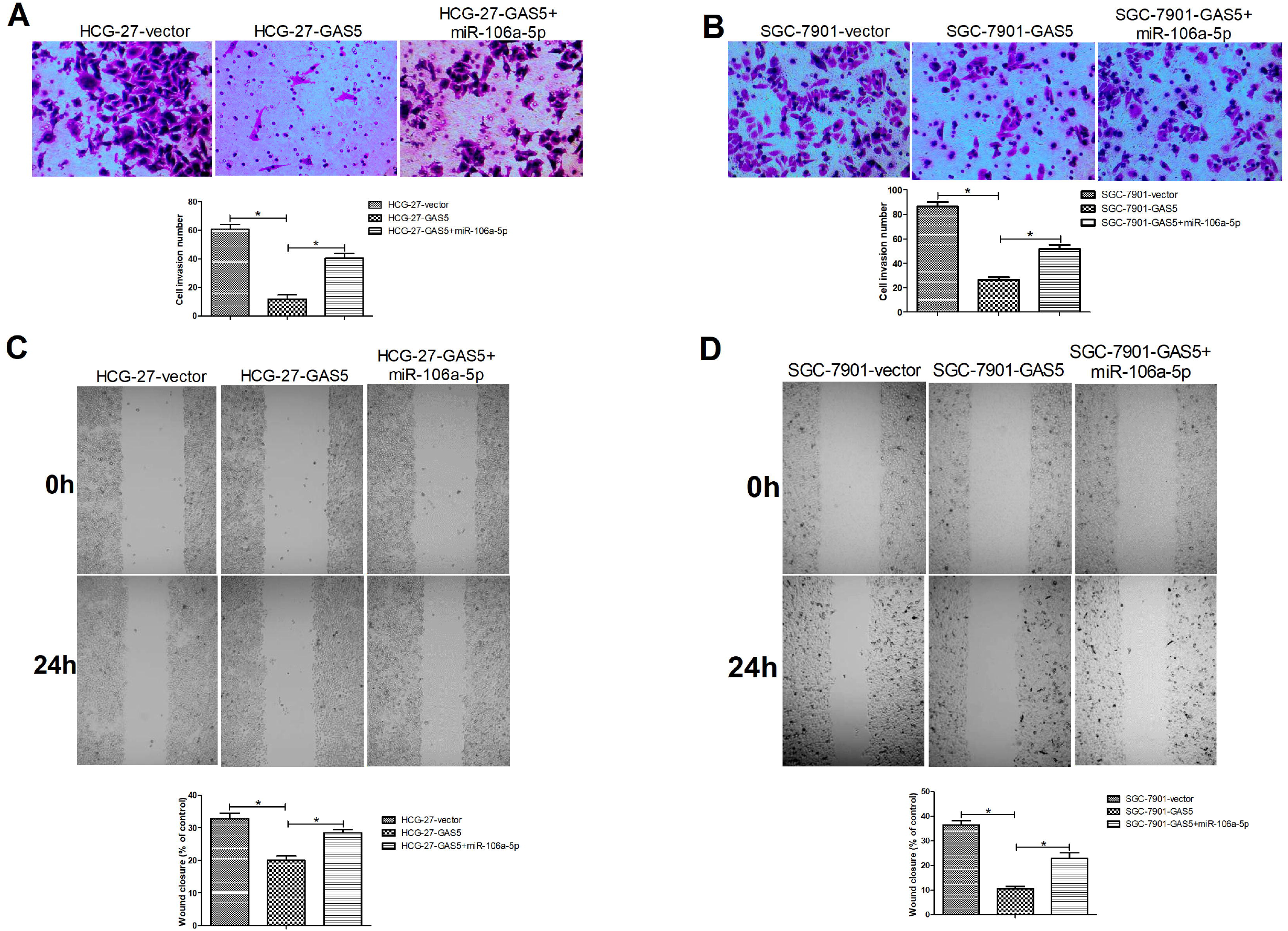
GAS5 overexpression repressed invasion and migration via regulating miR-106a-5p in GC cells. Transwell assay was carried out to assess the invasion capability of HCG-27 (A) and SGC-7901 (B) cells transfected with GAS5, GAS5 + miR-106a-5p, or vector at 0 and 24 h. And wound healing assay was used to determine the migration capability of the transfected HCG-27 (C) and SGC-7901 (D) cells. *\*P* < 0.05.

### GAS5 overexpression inhibited GC growth via regulating miR-106a-5p *in vivo*

To determine the function of GAS5 in tumor growth *in vivo*, the SGC-7901 cells transfected with GAS5, GAS5 + miRNA-106a-5p or vector were injected subcutaneously into nude mice. The detection of tumor size and weight showed that the volume of xenograft tumors with SGC-7901 cells expressing GAS5 was smaller than that of control group, and miRNA-106a-5p overexpression partially inversed the roles of GAS5 overexpression in GC tumor formation (Fig 5A and B). These results suggested that GAS5 overexpression inhibited growth of GC xenograft tumor via regulating miR-106a-5p.

**Figure 5.**
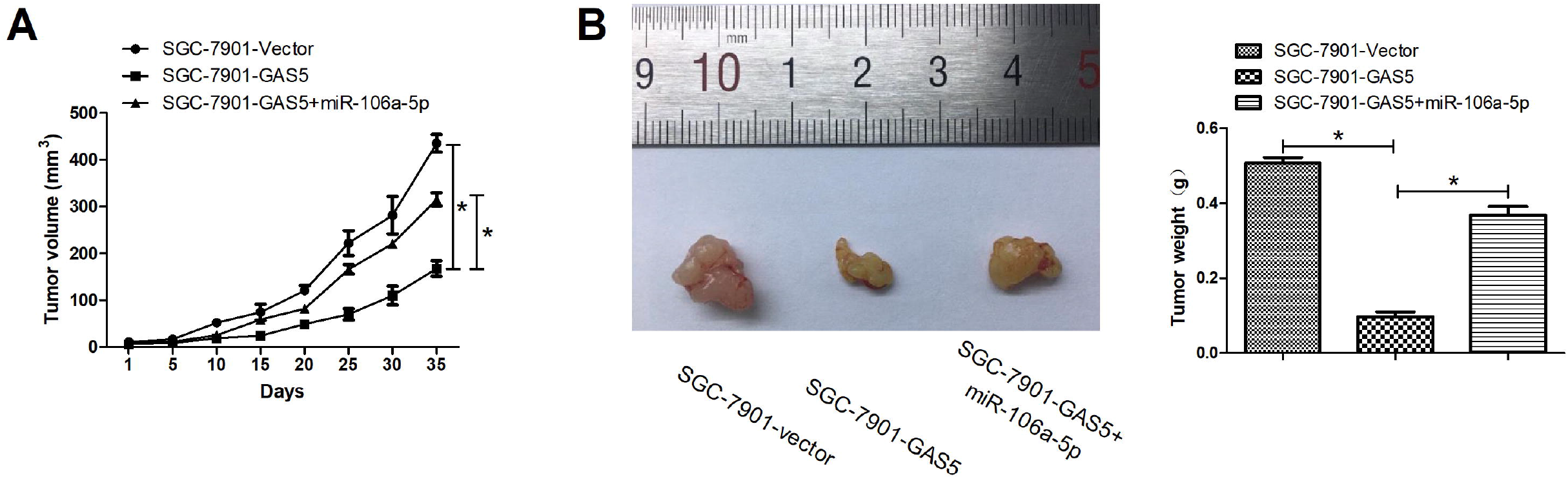
GAS5 overexpression inhibited GC growth via regulating miR-106a-5p *in vivo*. SGC-7901 cells transfected with GAS5, GAS5 + miR-106a-5p, or vector were subcutaneously injected into the right side of the posterior flank of mouse. Xenograft tumor growth curves (A) and tumor weights (B) for nude mice were detected. N = 7. Magnification, ×200. \**P* < 0.05.

**Figure 6.**
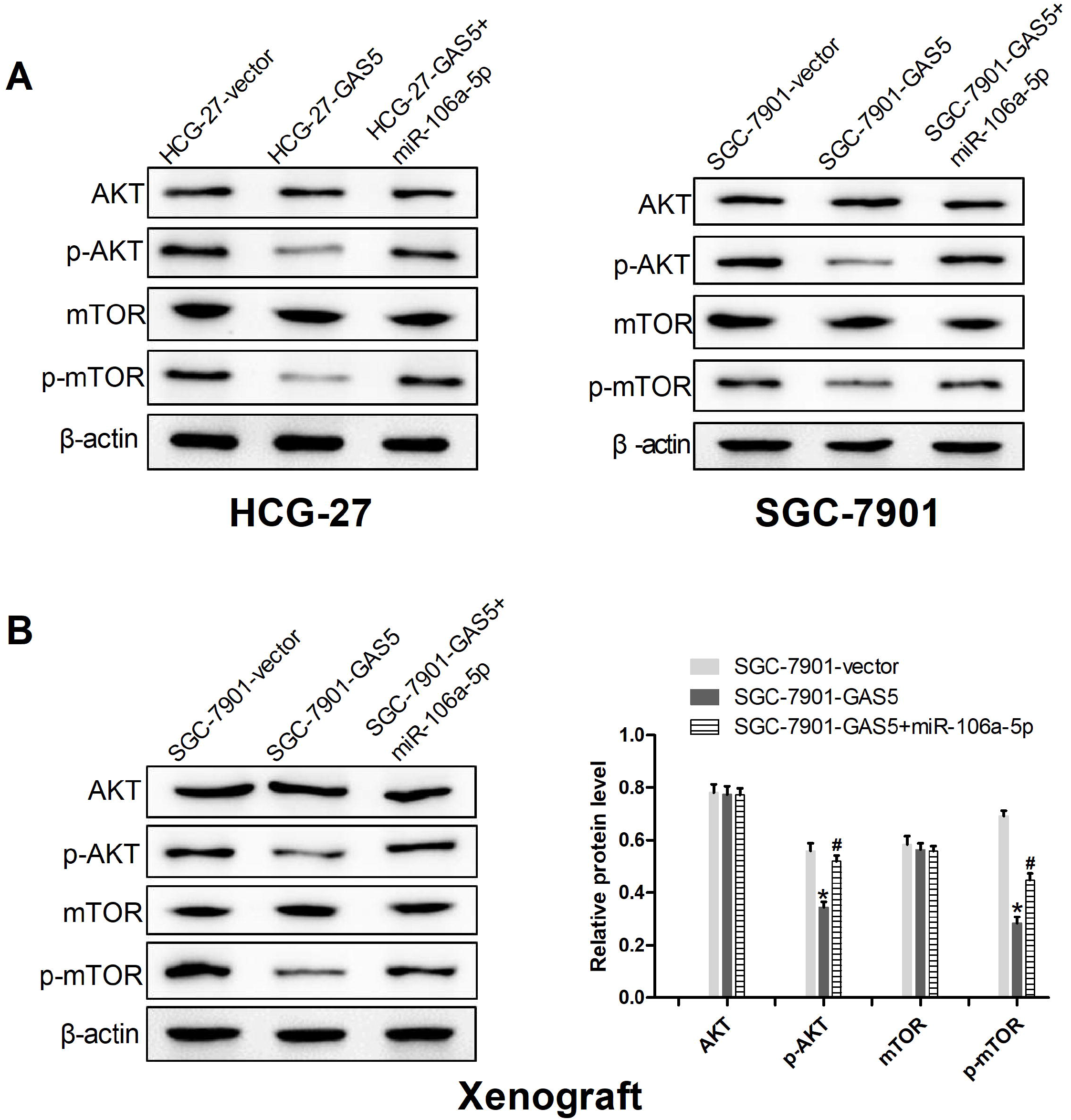
GAS5/miR-106a-5p axis negatively regulated the activation of the Akt/mToR pathway. (A) Western blot assay was conducted to detect the expression of Akt, p-Akt, mToR, and p-mToR in the HCG-27 and SGC-7901 cells transfected with GAS5, GAS5 + miR-106a-5p, or vector. (B) Western blot analyses of Akt, p-Akt, mToR, and p-mToR expression in xenograft tumor tissues. \**P* < 0.05: compared with vector-treated mice group, and ^#^*P* < 0.05: compared with the values of GAS5-treated mice.

### GAS5/miR-106a-5p axis negatively regulated the activation of the Akt/mToR pathway

The Akt/mToR pathway is an intricate intracellular pathway and its alterations are crucial in tumorigenesis (Cheng et al., 2015). To further explore the mechanism by which GAS5 and miRNA-106a-5p affected the GC progression, western blot assay was conducted to detect the effect of GAS5 and miRNA-106a-5p on activity of the Akt/mToR pathway *in vitro* and *in vivo*. The results from western blot exhibited that GAS5 upregulation significantly diminished the expression levels of p-Akt and p-mToR, while the introduction of miRNA-106a-5p partially inversed the repressive roles of GAS5 upregulation *in vitro* and *in vivo* (Fig A and B). These data showed roles of GAS5/miR-106a-5p axis in reducing activation of the Akt/mToR pathway in GC.

## Discussion

In the last decade, a growing number of cancer-related lncRNAs has been discovered; they are reported to be often abnormally expressed in numerous cancer cells and lead to carcinogenesis by spatially and temporally regulated expression patterns (Minotti et al., 2018). Although great progresses have been made in lncRNAs, their exact roles in GC are still not completely clear. GAS5, a lncRNA, was originally identified in growth-arrested mouse NIH 3T3 fibroblasts (Schneider et al., 1988). Afterwards, GAS5 has been found to be aberrantly expressed in many cancers and other diseases and played crucial roles in pathological process, such as cell proliferation, apoptosis, tumor growth and metastasis (Su et al., 2013). Pickard and his colleagues have presented abnormally low levels of GAS5 expression in prostate cancer cells and GAS5 upregulation boosts apoptosis and reduces 22Rv1 cell survival (Pickard et al., 2013). In pancreatic cancer, expression level of GAS5 has been verified to be obviously reduced in pancreatic cancer tissues and GAS5 upregulation represses cell proliferation by suppressing expression of cyclin-dependent kinase 6 (CDK6) *in vitro* and *in vivo* (Lu et al., 2013). In our study, we found that GAS5 expression was decreased in GC tissues and cells, and significant reduction of GAS5 levels in GC tissues at stage 3/4. Moreover, overexpression of GAS5 could restrain proliferation, invasion and migration as well as promote apoptosis in GC cells. These data revealed a crucial function of GAS5 in GC development.

miRNAs function vital regulatory roles in post-transcriptional expression of target gene and their aberrant expression are involved many human diseases including cancers (Li et al., 2018). Currently, some miRNAs have been reported to be related to GC. miR-25 has been reported to be upregulated in in plasma and primary tumor tissues of GC patients and enhances GC progression by directly targeting TOB1 (Li et al., 2015). Increased miR-34a could boost the sensitivity to DDP of SGC7901/DDP cells by suppressing cell proliferation and promoting cell apoptosis via targeting MET (Zhang et al., 2016). miRNA-106a, functioning as a cancer-promoting gene, has been shown to be correlated with carcinogenesis in many cancers, including GC (Yuan et al., 2016, Tian et al., 2018). In our study, our results showed that miRNA-106a-5p was significantly upregulated in GC tissues and cells. In recent years, growing number of reports have showed that the disorders of competitive endogenous RNA (ceRNA) networks tend to result in carcinogenesis by tumor suppressors and oncogenes through their ceRNA function (Cardoso et al., 2018, Liu et al., 2018, Zhang et al., 2018). Nonetheless, the ceRNA hypothesis has provided a novel viewpoint to explore disease processes by regulating ceRNA networks through microRNA (miRNA) competition or sponge for cancer therapy. In our results, what was interesting was that the expression of GAS5 is negatively correlated with miRNA-106a-5p expression. We speculated that GAS5 played important roles in GC progression by interacting with miRNA-106a-5p. As we had hypothesized, GAS5 directly targeted miR-106a-5p and negatively regulated its expression in GC cells. Furthermore, overexpression of miR-106a-5p could partialy inverse the effects of GC overexpression on cell proliferation, apoptosis, invasion, migration and xenograft tumor growth. All these result suggested that GAS5 could constrain GC progression.

miRNAs play vital roles in regulation of cell signaling and homeostasis. Plentiful of studies have showed that aberrations in various cellular signaling pathways are involved in regulating tumor development and progression (Polivka and Janku 2014). Mourtada et al have found that GAS5 upregulation could not result in growth arrest by itself, but enhance sensitivity to treatments that induce apoptosis through affecting diversified pathways (Mourtada-Maarabouni et al., 2009). Many reports have revealed that PI3K/Akt/mTOR, a main intracellular signaling pathway, has participated in cancer processes, such as cellular survival, proliferation, growth, invasiveness, and other functions of co-carcinogenic factor (Cheng *et al*. 2015, Li et al., 2015). As presented in our results, western bolt analysis showed that with the treatment of GAS5 transfection, the reducing phosphorylation of Akt and mTOR was determined, which was consistent with previous studies (Yacqub-Usman et al., 2015, Li et al., 2016). And miR-106a-5p overexpression could partialy inverse the repressive effects of GAS5 overexpression on activation of the Akt/mTOR pathway. Based on our results, we suggested that GAS5 inhibited tumorigenesis and development of GC by targeting miRNA-106a-5p through reduced the activation of the Akt/mTOR pathway.

## Conclusion

In conclusion, our present study revealed the decreased GAS5 and increased miR-106a-5p in GC tissues and cell lines. In addition, GAS5 competitively binds to miRNA-106a-5p and inhibited its expression in GC cells. Further, our results showed that GAS5 overexpression constrained cell proliferation, invasion, migration, and tumor growth as well as induced apoptosis by regulating miR-106a-5p through the Akt/mTOR pathway. Our findings provided a novel regulatory axis of GC progression, which may boost further exploration of functional lncRNA directed diagnostic and therapeutic agents in gastric cancer.

## Materials and methods

### Patients and tissue samples

The Fresh-frozen GC and paired adjacent normal epithelial tissue samples were obtained between from January 2017 to May 2018 from 21 GC patients who underwent surgery and didn’t receive any therapy in prior the operation at the First Affiliated Hospital of Zhengzhou University. Written informed consents were obtained from all the patients. The use of samples for all study was approved by the by the Research Ethics Committee of the First Affiliated Hospital of Zhengzhou University. All tissues were frozen in liquid nitrogen and then stored at −80°C for further qRT-PCR analyses.

### Cell culture

Human gastric epithelial cell line RGM-1 cell and gastric cancer cell lines (HCG-27 and SGC-7901) were used in this study and all cell lines were purchased from the Cell Bank of Type Culture Collection of Chinese Academy of Sciences (Shanghai, China). All cell lines were routinely grown at 37°C in an atmosphere of 5% CO_2_ in RPMI 1640 medium (Corning, Manassas, Virginia, USA) supplemented with 10% inactivated fetal bovine serum (FBS, Corning).

HEK293T cells were obtained from American Type Culture Collection (ATCC; Shanghai, China) and propagated in Dulbecco’s Modified Eagle’s Medium (Gibco; Invitrogen; Life Technologies, Germany) containing with 10% FBS under the above conditions.

### Gene overexpression

The complementary DNA (cDNA) encoding GAS5 or miRNA-106a-5p was PCR-amplified and subcloned into pCDH-CMV-MCS-EF1-copGFP-T2A-Puro vector (GenePharma, Shanghai, China). The recombinant vectors containing GAS5 or miRNA-106a-5p genes were co-transfected with pCMV-Δ8.2 and pCMV-VSV-G into HEK293T cells to generate retroviral particles overexpressing GAS5 or miRNA-106a-5p which were named as LV-GAS5 and LV-miR-106a-5p, respectively.

HCG-27 and SGC-7901 were transduced with GAS5 overexpressing lentivirus-containing media and screened with puromycin (2 μg/ml) to harvest the stable cells. After a week cell pools were obtained and expanded. miRNA-106a-5p overexpressing lentivirus was used to infect the GAS5 overexpressing HCG-27 and SGC-7901 cells to gain the cells with stable co-expression of GAS5 and miRNA-106a-5p.

### Luciferase reporter assay

The GAS5 fragment sequences holding the binding sites of miRNA-106a-5p were synthesized and then inserted into the pGL3 luciferase reporter vector (Promega, WI, USA). HEK293T cells were co-transfected with the constructs encompassing the wild type GAS5 (GAS5-WT) or mutant GAS5 (GAS5-MUT), along with the pRL-TK control vector and miRNA-106a-5p mimics (miRNA-106a-5p) or mimic control (miR-control). After48 h of transfection, the cells were collected and detected with Dual Luciferase Assay kit (Promega) in line with the manufacturer’s instructions.

### RNA extraction and quantitative real time PCR

Total RNA containing small RNA was prepared from either gastric cancer tissues or cells by TRIzol Reagent (Invitrogen; Thermo Fisher Scientific, Inc., Waltham, MA, USA) in according with the manufacturer’s instructions. Concentration and quality of RNA were measure using the NanoDrop ND-1000 spectrophotometer (NanoDrop Tech., Inc. Wilmington, DE, USA). Afterwards, 1 μg of RNA for GAS5 was used to synthesize cDNA using high Capacity cDNA Reverse Transcription Kit (Applied Biosystems, Foster City, CA, USA) in a T-Professional Basic, Biometra PCR System (Applied Biosystems). Reverse transcription of miRNA-106a-5p was performed to synthesize cDNA using TaqMan MicroRNA Reverse Transcription kit (Applied Biosystems). GAPDH and U6 as the endogenous controls were chosen to normalize the data of GAS5 and miRNA-106a-5p expression, respectively. All primers used in this study were shown in table 1.

**Table 1.**
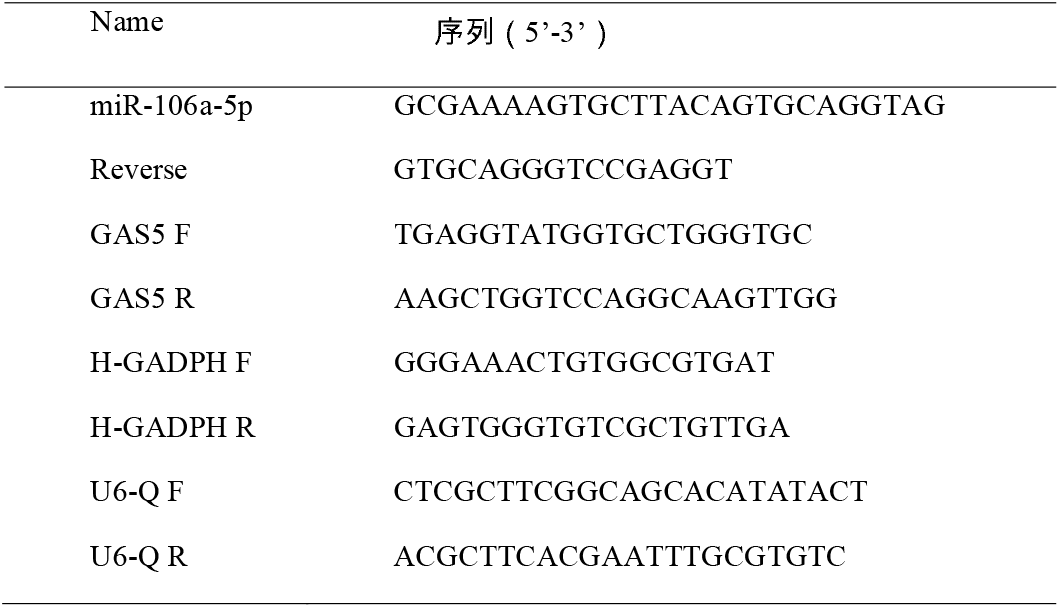
Primers used for quantitative real time PCR.

### CCK-8 assay

CCK-8 assays were used to assess the proliferation of cells. Briefly, the transfected cells at a density of 2×10^3^ cells/well were plated in 96-well plate. After incubation for the indicated times (0, 1, 2, 3 and 4 days), the CCK-8 solution (DOJINDO, Japan) was added into each well according the manufacture’s introduction. The optical density at 450 nm was determined by a microplate reader (Synergy4; BioTek Instruments, Inc., Winooski, VT, USA).

### Apoptosis analyses

Apoptosis rates of the transfected cells were detected by flow cytometric analyses. To measure the apoptosis rates in the transfected cells, the cells were grown for 24 h and then gathered with trypsin. For apoptosis analysis was performed by a double staining Annexin V and 7-AAD Apoptosis Detection Kit (BD Pharmingen, San Jose, CA, USA) following the manufacture’s introductions. Both cell cycle and apoptosis analyses were analyzed with a flow cytometer (FACScan; BD Biosciences).

### Wound healing assay

Cell migration capability was determined by wound healing assay. The transfected cells were plated in 24-well plate with complete RPMI 1640 medium. A 10-μl sterile pipette tip was used to make a vertical wound until cell confluence rate was about approximately 100%. The cells were washed with PBS to discard the detached cells and continuously cultured in FBS-free RPMI-1640 with for 24 □ h. Images were photographed at 0 and 24 h under an optical microscope (Olympus, Tokyo, Japan).

### Matrigel invasion assay

Cell invasion ability was assessed by matrigel invasion assay. The transfected cells were plated on the upper migration chambers (8 μm pore size; Millipore, Switzerland) coated with Matrigel (Sigma-Aldrich, USA) and incubated in free-FBS RPMI 1640 medium. 500 μL of RPMI 1640 medium supplemented with 20% FBS was added into the lower chamber. After incubation for 24 h, the invasive cells located on the lower surface of the chamber were stained with 0.05% crystal violet (Beyotime, Shanghai, China). The number of migrated cells was detected under an optical microscope (Olympus).

### *In vivo* animal studies

21 Four-week-old female BALB/c athymic nude mice were purchased from (the Experimental Animal Centre of Zhengzhou University) and maintained under specific pathogen-free conditions. Animal experiments were approved by the Animal Care and Use Committee of The First Affiliated Hospital of Zhengzhou University. The SGC-7901 cells transfected with GAS5, GAS5 + miRNA-106a-5p or vector were resuspended at a concentration of 5×10^7^ cells/ml. 200 μl of the suspended cells was subcutaneously injected into the right side of the posterior flank of each mouse. Tumor growth was monitored every 5 days, and tumor volumes were calculated using the equation V =π/6 × D × d2 (where V, volume; D, longitudinal diameter; d, latitudinal diameter). At 35 days post-injection, all mice were sacrificed, and weight of each tumor was detected. Tumor tissues were used for western blot analyses.

#### Western blot

Total proteins were extracted from tissue and the transfected cells using RIPA buffer (Beyotime) encompassing protease inhibitors. The protein lysates were separated by 10% SDS and then transferred onto a polyvinylidene fluoride membrane. The proteins were blocked with 5% non-fat milk in PBST (0.2% Triton X□100 in PBS) and the membranes were incubated with primary antibodies. The antibodies used in this study were as follow: AKT p-AKT, mTOR, p-mTOR, and β-actin, and all antibodies were purchased from Abcam Inc. (Cambridge, UK). After incubation with horseradish peroxidase (HRP)-conjugated secondary antibody for 1 h at room temperature, the immunoreactive proteins were signaled by using enhanced chemiluminescence (CEL; Thermo Fisher Scientific, Inc.). Finally, the protein expression levels were analyzed by ImageQuant software (LLC, Sunnyvale, CA, USA).

#### Statistical analysis

All data were shown as the mean ± standard deviation (SD). Statistical analyses were carried out by GraphPad Prism software. The Student’s t test was applied for comparisons between two groups and *P* < 0.05 was considered statistically significant.

### Competing interests

The authors declare no competing or financial interests.

### Author contributions

Conceptualization: S.D., X.Z.; Methodology: S.D., X.Z.; Software: D.L.; Validation: S.D., D.L.; Formal analysis: S.D., D.L.; Investigation: S.D., X.Z. Data curation: D.L.; Writing - original draft: S.D.; Project administration: S.D., X.Z.

